# Distinct contributions of memorability and object recognition to the representational goals of the macaque inferior temporal cortex

**DOI:** 10.1101/2025.10.06.680822

**Authors:** Soroush Ziaee, Ram Ahuja, Sabine Muzellec, Ezgi Fide, R. Shayna Rosenbaum, Kohitij Kar

## Abstract

The primate inferior temporal (IT) cortex, at the apex of the ventral visual stream, encodes information that supports diverse representational goals—from recognizing objects to determining which images are likely to be remembered. Specific artificial neural networks (ANNs), that currently serve as the leading computational hypotheses of ventral stream processing, are typically trained exclusively for object recognition. We asked whether incorporating image memorability as an additional optimization objective could improve ANN–brain alignment. Models optimized for memorability explained additional, non-overlapping variance in IT responses beyond that captured by recognition-optimized networks, indicating that memorability and recognition rely on partly independent dimensions of IT representation. Notably, these models also exhibited fewer non–brain-like units, bringing their representational geometry closer to that of IT. Furthermore, networks jointly optimized for both objectives were more predictive of human memorability than memorability-only models, while maintaining their alignment with human object recognition performance patterns. Together, these findings suggest that IT encodes multiple representational goals and that models trained solely for recognition provide an incomplete account of ventral stream computation.

**Significance:** Brain regions often serve multiple representational goals, and identifying those goals is critical because they provide the key to building better encoding models of the system. The primate ventral visual stream has traditionally been understood as a pathway for object recognition, with the inferior temporal (IT) cortex regarded as its core substrate. However, IT responses also predict image memorability—a robust phenomenon whereby some images are consistently remembered better than others. Here we show that memorability constitutes a separable representational goal of IT. ANNs optimized for memorability explained neural variance not captured by recognition models, and the two objectives produced distinct representational geometries. Critically, models jointly optimized for both recognition and memorability provided the best match to IT responses, improved prediction of human memorability, and preserved recognition performance. These findings highlight memorability as an organizing principle of IT and demonstrate that multi-goal optimization yields more brain-like computational models of vision.

## Introduction

What are the representational goals of the primate visual system? The visual system transforms complex sensory input into neural codes that guide a wide range of behaviors, from rapid recognition to long-term memory. Within this system, the ventral visual stream has long been recognized as the critical pathway supporting object recognition ^1,2^. Neurons in the inferior temporal (IT) cortex exhibit selectivity for object identity ^3,4^ and other visual forms ^5^, and lesion studies have established their necessity for object perception ^6–10^. For decades, object recognition has therefore been treated as one of the primary canonical computational goals of the ventral stream^11,12^. This perspective has also shaped the development of computational models, where artificial neural networks (ANNs) trained for large-scale object categorization tasks have emerged as the dominant framework for approximating IT responses and human behavior^13^.

Recognition-optimized ANNs, particularly deep convolutional networks trained on datasets such as ImageNet^14^, have been strikingly successful at predicting IT neural activity^15^. They explain a large, but partial fraction of the neural variance^12,13^, reproduce image-level patterns of object discriminability^16^, and capture the hierarchical representational transformations observed along the ventral stream^12,17^. These successes have reinforced the assumption that object recognition is not only the primary behavioral function of IT, but likely also a sufficient target for building accurate computational models of its responses. Yet IT representations are not limited to supporting object recognition alone. A growing body of evidence suggests that IT activity also aligns with other behavioral dimensions ^18–20^. One striking example is image memorability^21,22^ — the phenomenon that certain images are reliably remembered better than others^23^ across different tasks, contexts, and populations. Critically, this image-level property has been directly linked to IT responses: images that evoke stronger IT activity are more likely to be remembered later ^19^. These findings point to a broader role for IT, in which its representations are shaped not only by the demands of categorization but also by pressures related to learning and memory.

Considering memorability in the context of ventral stream^19,21,24^ computation highlights a tension in the field. On the one hand, computational models of IT are almost universally trained with recognition as their sole objective. On the other hand, the empirical data suggest that IT carries variance predictive of memorability. This mismatch raises an important question: do models trained exclusively for recognition omit critical dimensions of IT variance, and can incorporating memorability as an explicit training objective improve ANN–brain alignment? Addressing this question is significant for several reasons. First, it examines the representational goals of IT, inquiring whether memorability is encoded as a distinct dimension or as a byproduct of recognition that supports neural computations. Second, it tests a key assumption underlying current ANN–brain comparisons: that optimization for recognition alone is sufficient to explain IT. If memorability contributes unique variance, this assumption must be revised. Third, it introduces a more general principle: that building accurate models of high-level cortex may require multi-goal optimization^25,26^, reflecting evidence that biological systems are simultaneously shaped by multiple behavioral demands.

Memorability itself provides an especially compelling test case for multi-goal optimization. Unlike recognition, which depends on categorical distinctions between objects, memorability reflects the likelihood of encoding a particular image into long-term memory. The two goals overlap but are not identical: features that support recognition, such as shape or diagnostic parts, are not necessarily those that make an image memorable, whereas memorability may depend on distinctiveness, salience, or other factors that are not critical to recognizing an object. Behaviorally, this dissociation is clear: images can be easily recognized but poorly remembered, and vice versa. For example, the make and model of a car provide features that support recognition, allowing it to be categorized as a car regardless of context. In contrast, a broken window or unusual detail might not aid recognition, but it may make the image more distinctive and thus more likely to be remembered. Neurally, the same IT responses that predict recognition accuracy ^27,28^ also predict memorability^19^, but not perfectly. Thus, recognition and memorability appear to represent partly independent dimensions of IT coding.

If both object recognition and image memorability constitute functional dimensions of IT coding, then models optimized solely for recognition are expected to capture the variance associated with categorical discrimination while failing to fully account for variance uniquely linked to memorability. Incorporating memorability as an explicit optimization objective should therefore enhance the fidelity with which artificial neural networks approximate IT responses.

To directly test this idea, we trained ANNs under three regimes—recognition-only, memorability-only, and joint recognition-plus-memorability—and evaluated them against macaque IT responses, human memorability judgments, and human and monkey recognition behavior. We found that memorability optimization explained complementary IT variance beyond recognition, that the two objectives drove models toward distinct representational geometries, and that joint training improved alignment with both IT and human memorability while preserving recognition performance. Together, these results demonstrate that IT encodes multiple representational goals and underscore the need for multi-goal optimization to build models that more faithfully capture the richness of biological vision.

## Results

To test whether incorporating memorability as an explicit optimization objective improves the ability of artificial neural networks (ANNs) to model responses in inferior temporal (IT) cortex, we conducted a series of analyses that proceeded in four stages. We first characterized human behavior on object recognition and image memorability tasks and then examined whether macaque IT responses capture these behavioral patterns. IT population activity predicted image-level variation in both recognition accuracy and memorability, confirming that IT encodes variance aligned with both behavioral goals. We next compared the representational similarity of recognition-versus memorability-optimized models to determine whether these objectives yield distinct internal feature geometries. We then evaluated how well each model family predicts IT responses and whether the variance explained by recognition and memorability is shared or unique. Finally, we tested whether jointly optimizing for both objectives produces models that better capture IT responses and behavioral outcomes than recognition-only networks.

### Object discriminability for images do not fully explain its memorability

We first asked whether image memorability could be fully explained by object discriminability. Human participants (n= 88) performed a binary object discrimination task across 50 naturalistic images drawn from 10 categories (5 images per object category). On each trial, a test image was shown for 100 ms, followed by two canonical images: one of the target object and one of a distractor (Figure 1A). Distractors were drawn from all nine non-target categories, and each image was tested against every possible distractor. We then computed the image-level recognition accuracy as the average performance across all comparisons. In a separate task with the same images, participants (n = 22) viewed each stimulus for 100 ms and judged whether it was novel or repeated after a fixed lag of 10 intervening fillers (Figure 1B). From these responses we derived image-level memorability scores, reflecting the probability of correct recognition upon repetition. Across images, object discriminability and memorability were only weakly correlated (Pearson *r(48)* = 0.18, *p* = 0.20; Figure 1C), indicating that the ability to recognize an object category does not fully predict which specific images are more likely to be remembered. This dissociation suggests that image memorability reflects an additional representational dimension beyond object-level discriminability.

**Figure 1.**
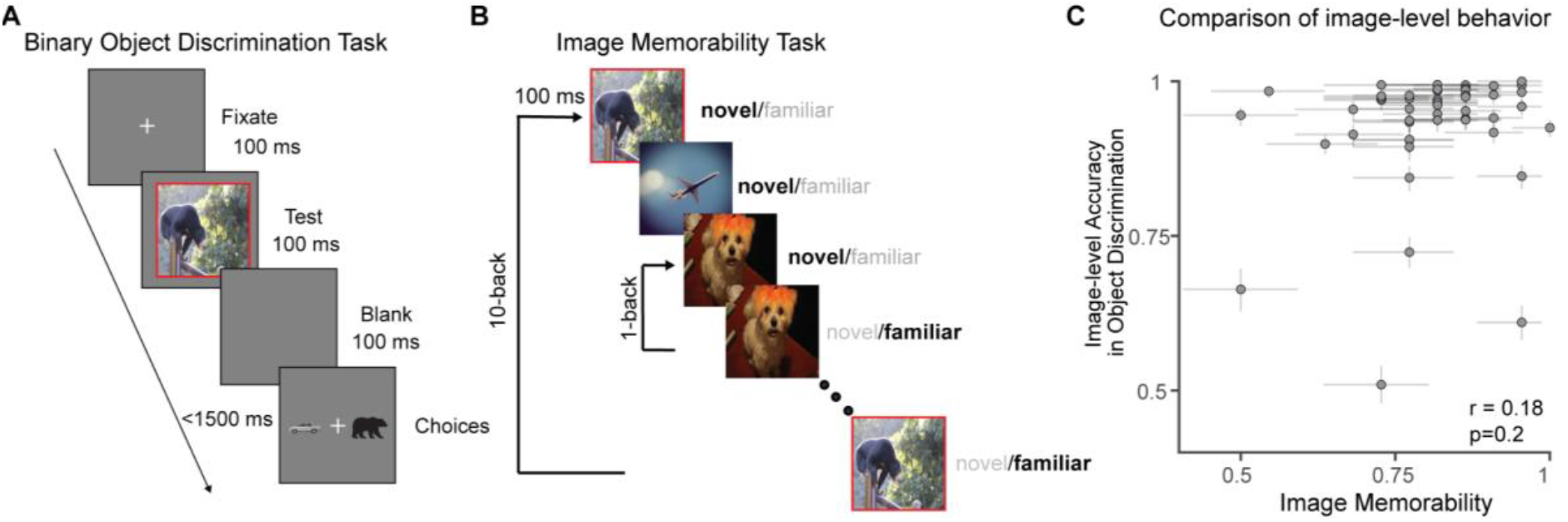
Object discriminability for images does not fully explain its memorability. **A**. *Binary object discrimination task*. After fixation (100 ms), a test image (100 ms) was presented, followed by a 100 ms blank and a two-alternative choice screen (<1500 ms) with a canonical target and a canonical distractor object. Each image was tested against all nine non-target categories to obtain image-level accuracy. **B**. *Image memorability task*. Each image (100 ms) appeared as novel or was repeated after a lag of 10 intervening images; participants reported whether each image was novel or familiar on every trial. A few trials served as “catch” trials that were repeated immediately (1-back, lag = 0) to monitor attention and reject inattentive subjects. **C**. *Comparison of image-level behavior*. Scatterplot comparing human memorability scores (x-axis) and image-level accuracy in the human object discrimination task (y-axis), across 50 images. The weak correlation (*r* = 0.18, *p* = 0.2) indicates that object discriminability does not fully account for image memorability, suggesting that memorability reflects a separable dimension of visual processing beyond object recognition. Error bars indicate 95% confidence intervals of estimating the image memorability and object discrimination accuracies.

### IT responses predict both recognition and memorability behavior

The macaque IT cortex has long been regarded as the neural substrate of object recognition^1^. Decades of neurophysiology and computational modeling have demonstrated that IT activity closely tracks object identity and predicts behavioral performance on recognition tasks^28^. To verify this, we measured large-scale neural responses across the IT cortex of two monkeys (Figure 2A, 155 sites in monkey 1, 109 sites in monkey 2; see inclusion criteria). Monkeys passively fixated images (1320 images) shown in their central 8° of visual field for 100 ms, randomly interleaved, with a gray blank screen for 100 ms in between each presentation (Figure 2A). We then decoded object identity from these IT responses by training cross-validated linear one-vs-rest classifiers on population activity measured 70–170 ms after image onset (as in previous studies^27–29^). The outputs of these classifiers were used to generate image-level predictions of recognition behavior at the image-level for the 50 images used in the behavioral study (see Methods). Across images, IT activity robustly predicted recognition performance, replicating the canonical link between IT and categorization accuracy. Because electrophysiological recordings necessarily sample only a small fraction of the neurons in IT, we also examined how the strength of the neural–behavioral relationship depends on population size (Figure 2B). We next asked whether the same IT responses also predicted image memorability. To quantify how well neural activity in inferior temporal (IT) cortex predicts human image memorability, we computed the correlation between population response magnitude and behavioral memorability scores across images. For each image, the population response magnitude was defined as the Euclidean (L2) norm of the mean response vector across simultaneously recorded neurons, representing the overall strength of IT activation to that image. Consistent with prior reports^19^, images that evoked stronger IT responses were also more likely to be remembered (high predictivity ∼0.5 of memorability, as neural sampling size increases: Figure 2C). This relationship was robust across images and aligned with the broader evidence that memorability is a reliable and generalizable correlate of IT (Figure 2C). Across images (n = 50), the accuracy of IT-based object discrimination decodes was not correlated with mean population response magnitude (r = –0.04, p = 0.73), indicating that overall response strength does not trivially account for image-level object discriminability (Figure 2D). Together, these analyses establish that IT representations align with both recognition and memorability, motivating the question of whether artificial neural networks trained solely on recognition objectives can capture this dual representational role, or whether memorability contributes complementary variance in IT.

**Figure 2.**
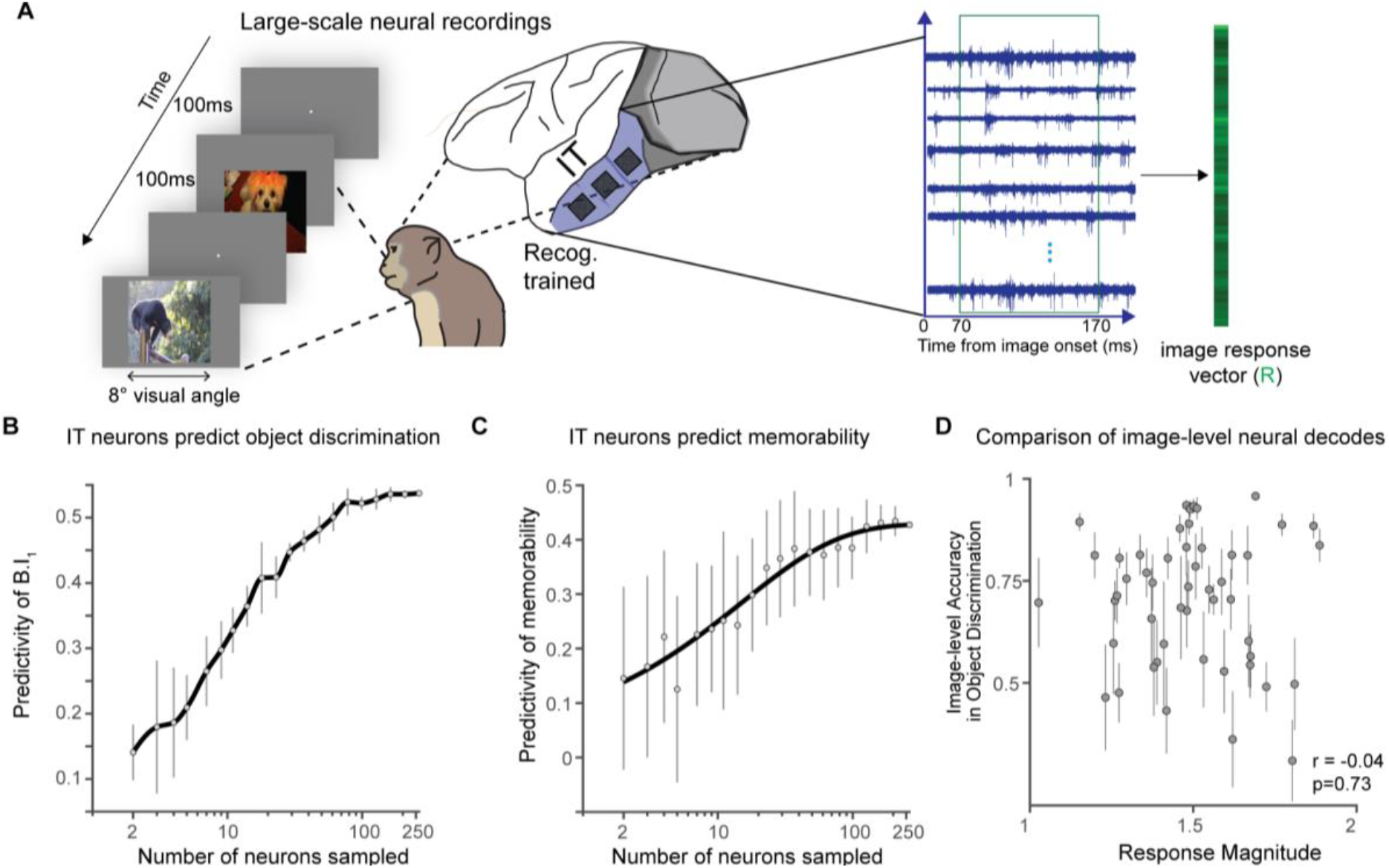
Distributed neural activity in the macaque IT cortex predicts both image memorability and object discrimination. **A**. Large-scale electrophysiological responses were recorded from macaque IT cortices during rapid image presentations (100 ms) at an 8° visual angle. Population response vectors (*R*) were extracted from 70–170 ms after image onset. **B**. *IT predicts object discrimination*. The predictivity of the image-level one-versus-all metric B.I_1_ from IT responses increases and then saturates with population size (circles: mean across bootstrap subsamples; vertical bars: variability; black line: smoothed trend). **C**. *IT predicts memorability*. Predictivity of human memorability from IT population responses increases with the number of randomly sampled IT neurons (format as in B). Error bars in B and C indicate the standard deviations across neural sampling. **D**. *Comparison of image-level neural decodes*. Across images (n=50), the accuracy of IT-based object discrimination decodes was not correlated with mean response magnitude (*r* = –0.04, *p* = 0.73). Error bars indicate the standard deviation across cross validation splits.

### Recognition- and memorability-optimized models develop distinct representational spaces

We next asked whether training ANNs to optimize for memorability (with the LaMem dataset^30^) produces representational geometries similar to, or distinct from, those optimized for object recognition (with the ImageNet dataset^14^). To address this, we trained (see Methods) 26 ANN models (base architectures, and IT-like layers shown in *Table 1*), on these two objectives respectively, and then compared (Figure 3A) their internal feature spaces for the IT-like layers (as determined from Brain-Score^13^) using centered kernel alignment (CKA^31^). We used CKA to quantify similarity in representational structure across network layers. Within the same architecture, recognition- and memorability-trained networks diverged substantially, with the largest differences emerging in deeper layers (Figure 3B). This result indicates that the optimization objective, rather than the architecture alone, plays a central role in shaping representational geometry. We further compared representational similarity between models trained on true memorability scores and those trained on ImageNet labels using CKA (Figure 3C). Across architectures, memorability-optimized models showed consistently lower CKA similarity with recognition-trained networks, in their IT-like layers, indicating that optimization for memorability drives a systematic representational shift relative to recognition. Together, these findings demonstrate that recognition and memorability optimization lead ANNs toward distinct representational spaces. This provides a mechanistic explanation for why recognition- and memorability-trained models might capture partly non-overlapping variance in IT responses.

**Table 1.** Base architectures and ANNs used in the study and the corresponding IT layers.

|  | ANN Model Name | IT-aligned Layer Name |
| --- | --- | --- |
| 1. | AlexNet<sup>55</sup> | features.12 |
| 2. | ResNet-18<sup>56</sup> | layer4.0.bn1 |
| 3 | ResNet-50<sup>56</sup> | layer3 |
| 4 | ResNet-101<sup>56</sup> | layer4.0.relu |
| 5 | Vgg16<sup>57</sup> | feature.23 |
| 6 | Vgg19<sup>57</sup> | feature.27 |
| 7 | Inception<sup>58</sup> | Mixed_7a |
| 8 | vit_b_16<sup>59</sup> | transformer.blocks.3.pwff.fc2 |
| 9 | vit_b_32<sup>59</sup> | transformer.blocks.4.pwff.fc2 |
| 10 | Swin_t<sup>60</sup> | trunk.layers.2.blocks.2 |
| 11 | Swin_b<sup>60</sup> | trunk.layers.2.blocks.4 |
| 12 | Swin_s<sup>60</sup> | trunk.layers.2.blocks.6 |
| 13 | squeezenet1_1<sup>61</sup> | features.9.expand3x3 |
| 14 | shufflenet_v2_x1_0<sup>62</sup> | stage4 |
| 15 | resnet50_swsl | layer3_fake_relu |
| 16 | mobilenet_v2<sup>63</sup> | mobilenet_v2.layer.12 |
| 17 | MnasNet | cell_12 |
| 18 | Efficientnet_v2_s<sup>64</sup> | features.7 |
| 19 | efficientnet_v2_m<sup>64</sup> | features.8 |
| 20 | efficientnet_b6<sup>65</sup> | features.8 |
| 21 | efficientnet_b4 | features.8 |
| 22 | densenet161<sup>66</sup> | features.transition3.pool |
| 23 | convnext_tiny | features.5.4.block.0 |
| 24 | convnext_small<sup>67</sup> | features.5.9.block.0 |
| 25 | convnext_large<sup>67</sup> | features.5.11.block.0 |
| 26 | convnext_base<sup>67</sup> | features.5.11.block.0 |

**Figure 3.**
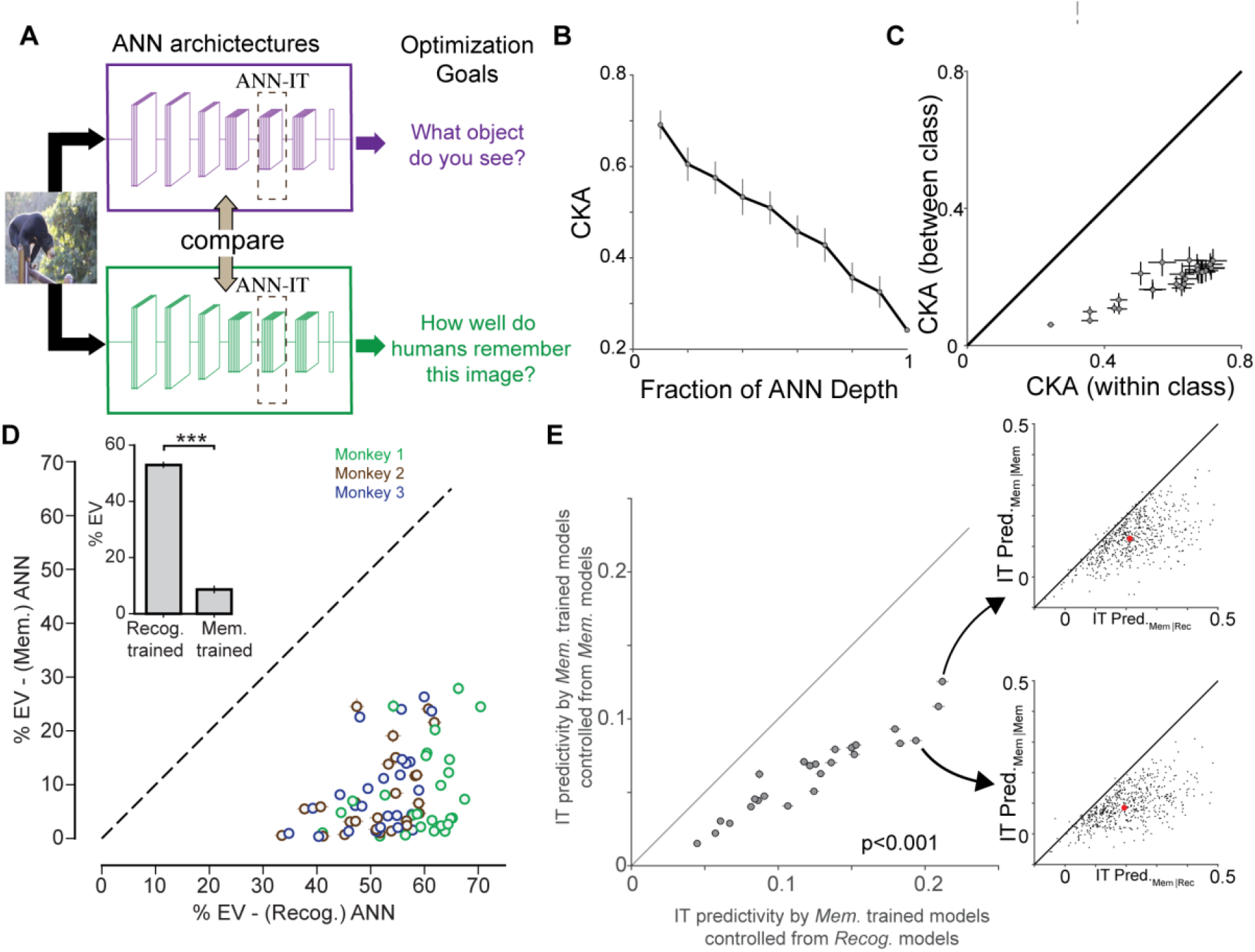
Recognition- and memorability-optimized models develop distinct representational geometries, predict IT responses but capture partly distinct variances. **A**. Schematic of the comparison: identical ANN architectures were trained with different optimization goals—object recognition (purple) or image memorability prediction (green)—and their internal feature spaces were aligned to IT cortex responses. **B**. Layer-wise representational similarity, measured with centered kernel alignment (CKA), between recognition- and memorability-optimized networks within the same architecture. Divergence increased with model depth, indicating that the optimization objective, rather than architecture alone, shapes deeper representations. **C**. Control analysis: CKA similarity between models trained with memorability scores (within class) versus with ImageNet labels (between class). Memorability training led to consistent representational separation from recognition-trained models. **D**. Predictivity of recognition versus memorability-trained models across images. Both model classes predicted IT responses significantly above chance (inset: group means with error bars, p < 0.01), but recognition-trained models explained more variance overall. **E**. *Cross-validated variance-partitioning analysis of IT predictivity*. Each point represents one model, showing its average ability to predict IT responses after controlling either for recognition model predictions (x-axis) or for a held-out trial set of its own memorability predictions (cross-validated self-control; y-axis). Points lying below the unity line indicate that memorability-trained models predict IT activity more strongly when controlling for recognition than when controlling for their own predictions, demonstrating that the additional variance captured by memorability models reflects genuine, behaviorally meaningful structure rather than noise or overfitting. Error bars denote ± 1 s.e.m. across neurons. Insets on the right show neuron-by-neuron data for two representative models, confirming the same pattern at single-cell resolution.

### Recognition and memorability models both predict IT responses, but account for partly distinct variance

We next quantified how well recognition- and memorability-optimized models predict macaque IT responses. For this analysis, we measured distributed neural responses across the macaque IT cortex of three macaques (Figure S3), with chronically implanted Utah arrays. We identified 571 visually driven IT sites while the three monkeys viewed 640 naturalistic images (previously used in^32,33^) presented for 100 ms at ∼8° visual angle. ANN features were linearly mapped to IT responses using ridge regression with cross-validation, and predictive accuracy was expressed as the percentage of explainable variance (%EV) on held-out images. Across architectures, recognition-optimized models consistently explained more variance than memorability-optimized models (Figure 3D, one-sided Wilcoxon signed-rank test: *z = 351, p < 0*.*001* for all three monkeys). At the population level, recognition models achieved mean %EV values in the range of 25–30%, with some neurons reaching nearly 60%, whereas memorability models averaged closer to 15–20% EV. Both sets of models nevertheless performed significantly above zero (one-sample *t*-tests, *p* < 0.01 for both), demonstrating that optimization for either behavioral goal yields features aligned with IT. To test whether the variance explained by memorability-trained models reflects true memorability structure rather than overfitting or shared variance with recognition, we performed a cross-validated variance-partitioning analysis (Figure 3E). On the x-axis, we plot the predictivity of IT responses by memorability-trained models while controlling for the recognition model’s predictions. On the y-axis, we plot the same measure but controlling for a held-out trial set of the same image responses predicted by the memorability model itself (cross-validated self-control). Each point represents one model, with values averaged across neurons (error bars: s.e.m. across neurons). The significantly higher x-axis values (paired t-test, t(25) = 12.81, p<0.001) indicate that residuals from the memorability model retain IT-predictive information beyond what is explained by recognition, whereas residuals from the cross-validated self-regression do not— confirming that the additional variance captured by memorability-trained models is genuine and not noise. The insets on the right show neuron-by-neuron results for two representative models, illustrating the same pattern at single-cell resolution. Together, these analyses demonstrate that memorability optimization captures a modest but detectable IT signal, while recognition accounts for substantially more unique variance. Although memorability is insufficient on its own to produce high-fidelity IT models, it contributes complementary structure absent from recognition-only training, providing a mechanistic explanation for why jointly optimized models could yield improved ANN–IT alignment.

### Joint optimization for recognition and memorability improves ANN–IT forward predictivity

We then asked whether combining recognition and memorability training could yield models that better approximate IT responses. Starting from ImageNet-pretrained networks, we jointly optimized each architecture using a Learning-without-Forgetting (LwF, see Methods) distillation protocol^34^, which retained recognition performance while learning image memorability. These jointly trained models outperformed recognition-only networks, capturing significantly higher explainable variance in IT responses across layers and architectures (one-sided paired tests, M1: *Z = 65, p = 0*.*002*, M2: *t(25) = -2*.*941, p = 0*.*003*, M3: *t(25) = -2*.*830, p = 0*.*005*)). Importantly, this improvement depended on the true memorability labels, and not simply on the exposure to more images (from the LaMem dataset). To verify this, we repeated the joint training procedure with shuffled memorability labels (Figure S1). The % EV of recognition+memorability models were significantly higher than recognition + memorability (shuffled) models (one-sided paired tests, M1: *Z = 64, p = 0*.*002*, M2: *Z = 82, p = 0*.*008*, M3: *Z = 126, p = 0*.*109*). Importantly, the architectures benefiting more in terms of IT predictivity from the joint optimization were also more affected by the shuffling (correlation of deltas: *r(25) = 0*.*57, p = 0*.*002*). Thus, the gains were not due to additional training exposure, but specifically to learning the memorability structure within the images. Together, these results provide direct evidence that models trained with multiple behavioral objectives better capture IT responses than recognition-only models. This supports our claim that IT is shaped by separable representational goals, and that models incorporating both recognition and memorability provide a more complete account of ventral stream computation.

To explore what factors determine the success of joint optimization, we next examined how gains in IT predictivity (%ΔEV) related to the behavioral and training properties of each model (Figure S2). Models that achieved higher memorability prediction scores exhibited greater increases in IT explained variance (*r* = 0.43, *p* = 0.02), indicating that improvements in ANN– IT alignment tend to scale with the fidelity of memorability learning. In contrast, the relationship between gains in %EV and changes in ImageNet classification accuracy followed a non-linear trend: networks that retained moderate recognition performance showed the largest increases in IT predictivity, whereas models with minimal or large drops in ImageNet accuracy showed little to no gain. This pattern suggests that successful joint optimization depends on achieving a balance between preserving recognition capability and learning memorability structure, mirroring the integration of multiple representational goals observed in the IT cortex.

### Joint optimization for recognition and memorability improves ANN–IT reverse predictivity

Thus far, our analyses have focused on forward predictivity—how well model features can be mapped onto IT responses. We next asked whether joint training also improves reverse predictivity, defined as the ability to reconstruct model feature space directly from IT neural responses (Figure 5A)^35^. Reverse predictivity provides a complementary measure of alignment, since models can differ in how well their representational geometry is embedded in neural activity even when forward predictivity is comparable. Consistent with previous work, we observed that forward and reverse predictivity were not strongly correlated across models (Figure S4), suggesting that the two metrics capture distinct aspects of brain–model alignment.

**Figure 4.**
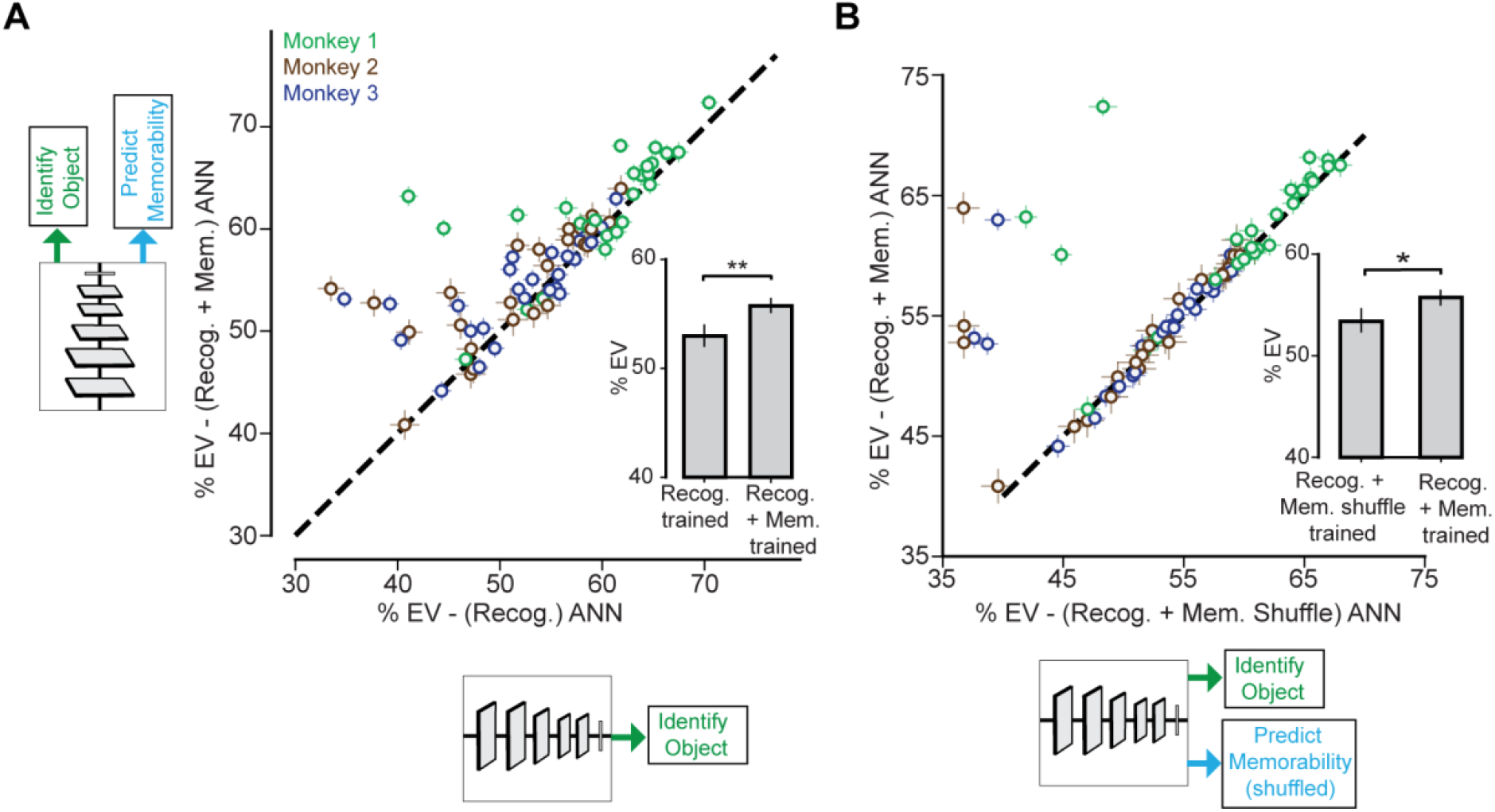
Joint optimization for recognition and memorability improves ANN–IT alignment. **A**. Schematic of network training and neural predictivity comparison between recognition-only models (x-axis), and recognition+memorability models (y-axis). Jointly optimized models explained significantly more IT variance than recognition-only models (inset: group means, *p < 0*.*05*). **B**. Comparison of recognition and memorability (shuffled) versus recognition and memorability (non-shuffled) models. Shuffled-label training did not improve predictivity over recognition-only baselines, confirming that gains required true memorability structure (inset: group means, *p < 0*.*05*).

**Figure 5.**
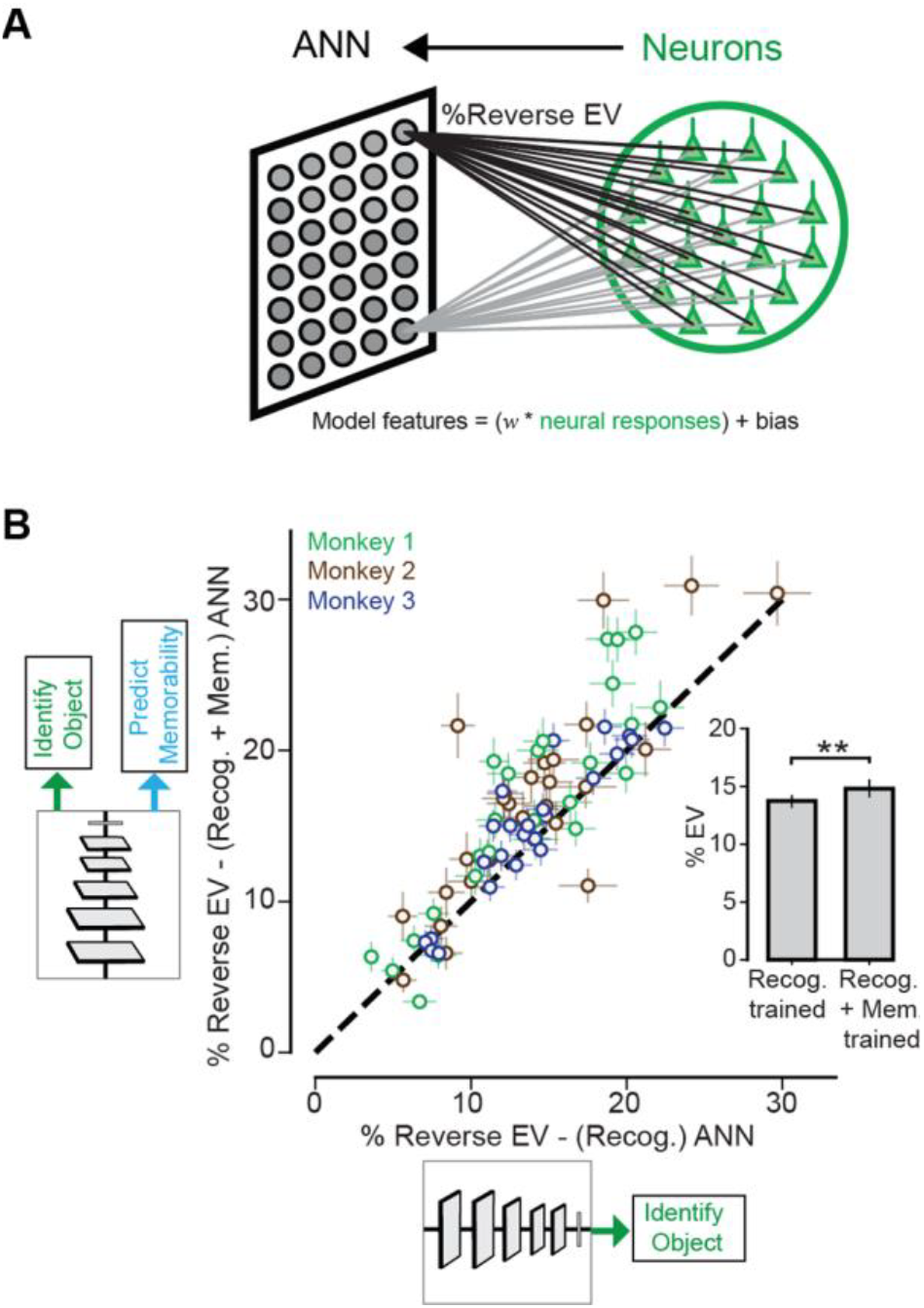
Joint optimization for recognition and memorability improves reverse predictivity. **A**. Schematic of reverse predictivity analysis: IT population responses were linearly mapped back into model feature space, and performance was quantified as the percentage of explained variance (% Reverse EV). **B**. Comparison of reverse predictivity for recognition-only versus recognition+memorability models. Jointly trained models (green/blue) consistently achieved higher % Reverse EV than recognition-only baselines (brown), with most points lying above the unity line. Inset:Group means across models confirmed a significant increase in reverse predictivity for jointly optimized networks (**p < 0.01**).

**Figure 6.**
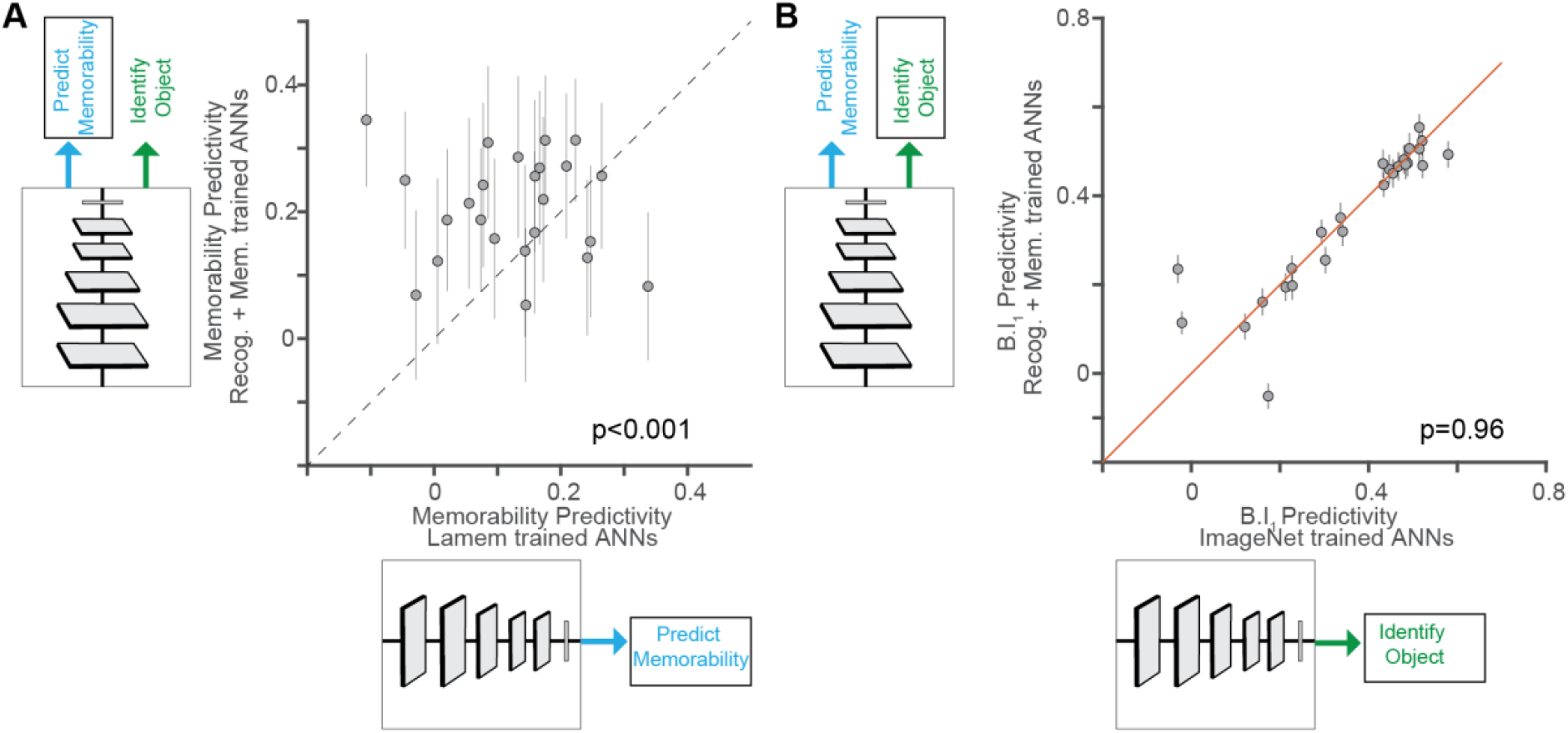
Jointly optimized models improve memorability prediction without altering recognition alignment. **A**. Comparison of memorability prediction between memorability-only ANNs and recognition+memorability ANNs. Jointly optimized models predicted human memorability scores significantly better than memorability-only models (p < 0.001). **B**. Comparison of recognition prediction (image-level B.I_1_ metric) between recognition-only ANNs and recognition+memorability ANNs. Performance was nearly identical (p = 0.96), indicating that joint optimization preserved recognition alignment.

Across architectures, recognition-only models achieved modest reverse predictivity, with mean explained variance (%EV) values of ∼12–14%. Jointly optimized recognition+memorability models yielded significantly higher reverse predictivity (mean %EV increase ≈ 2–3%, one-sided paired t-test, p < 0.01, M1: *t(25) = -3*.*99, p < 0*.*001*, M2: *t(25) = -3*.*59, p < 0*.*001*, M3: *t(25) = -3*.*13, p = 0*.*002*). At the single-neuron level, this improvement was consistent, with most points lying above the unity line when comparing recognition-only versus recognition+memorability models. Importantly, forward and reverse gains were not redundant: models that improved in forward predictivity did not necessarily improve in reverse predictivity (*r(25) = 0*.*3, p = 136*), yet both metrics increased under joint optimization.

These findings indicate that adding memorability as an explicit objective yields model representations that are not only better predictors of IT responses but are also better predicted from IT activity. This reciprocal alignment suggests that multiple behavioral constraints push models toward representational geometries that more closely match those instantiated in the brain.

### Jointly optimized models improve memorability prediction without altering recognition alignment

Finally, we asked whether the variance captured by joint training translated into better predictions of behavior. For memorability, we observed that jointly optimized models substantially outperformed memorability-only networks in predicting human image memorability scores (paired t-test; t(25) = -2.3968; p =0.02). This shows that the variance gained through joint training directly enhanced alignment with human memorability judgments. In contrast, jointly optimized models did not provide additional benefits for object recognition behavior. Their predictions of human and monkey recognition accuracy were comparable to those of recognition-only models (paired t-test; t(25) = -0.0393; p = 0.96), indicating that the added variance was specific to memorability and did not contribute further to recognition alignment. This dissociation reinforces the view that recognition and memorability reflect partly separable dimensions of IT representation. Incorporating memorability into model training thus improves ANN–IT alignment for memorability-related variance while preserving recognition performance, underscoring the role of multiple optimization pressures in shaping IT representations.

## Discussion

Our study demonstrates that the macaque IT cortex encodes multiple representational goals, extending beyond its well-established role in object recognition. By showing that IT responses also predict image memorability, and that models trained jointly on recognition and memorability objectives align more closely with IT than recognition-only networks, our results provide direct evidence that ventral stream representations are shaped by separable optimization pressures. This finding challenges the long-standing assumption that recognition is the sole functional target of IT and suggests that memorability serves as a complementary organizing principle in the high-level visual cortex.

### Multiple representational goals in IT

For decades, object recognition has served as the dominant explanatory framework for IT function. Neurophysiological studies have established that IT responses carry sufficient information to support recognition behavior, and recognition-optimized ANNs have achieved considerable success as computational models of IT^2^. Yet this single-goal perspective has always been an incomplete account of IT variance: a substantial portion of neural responses remains unexplained by recognition alone. Our results identify memorability as one source of this variance.

Image memorability is a robust behavioral phenomenon that is consistent across observers. Previous work has linked memorability to IT response magnitude^19^, but whether this variance is functionally relevant and computationally useful has remained unclear^21^. Here, we demonstrate that memorability-optimized models capture non-overlapping variance in IT responses, and that joint optimization for memorability and recognition enhances ANN–brain alignment. These findings suggest that IT representations might be optimized not only to support rapid categorization of objects but also to prioritize which images are likely to be remembered.

A concrete way to illustrate this distinction is to again consider the car with a broken window. Features such as the size, make, or shape of the car are crucial for recognizing it as a car and discriminating it from other categories, like dogs or airplanes. In contrast, the broken glass does not improve categorical recognition but makes the image more distinctive and thus more memorable. Our results reflect exactly this kind of separation: recognition-optimized models explain most of the variance associated with categorical discrimination, while memorability-optimized models account for a smaller but complementary portion of variance related to what stands out and is retained in memory.

### Implications for computational models of vision

The discovery that memorability training improves ANN–IT alignment carries important implications for model development. Most ANN models of vision are trained on large-scale recognition benchmarks such as ImageNet^13,14^ and are evaluated primarily on recognition accuracy^16,36,37^. While this has driven remarkable engineering progress, it has also reinforced the implicit assumption that recognition is the primary optimization pressure shaping IT representation. Our results challenge this assumption. By demonstrating that recognition-only training leaves significant IT variance unexplained and that multi-goal training yields better alignment, we highlight the limitations of single-objective optimization. The finding that memorability-optimized models develop representational geometries distinct from those of recognition-optimized models underscores that optimization targets strongly influence the learned representations^25^. The success of joint training further suggests that ANNs can integrate multiple behavioral objectives^38^ without sacrificing performance on either, yielding richer and more brain-like feature spaces. This perspective aligns with recent calls to move beyond single-task benchmarks in computational neuroscience and toward models that reflect the diversity of behavioral goals guiding biological vision^39,40^.

### Role of the IT cortex in image memorability

One key interpretive question is why does IT activity correlate with the measures of image memorability in the first place. One possibility is that image memorability reflects a byproduct of IT selectivity: images that drive strong responses in a broad population of IT neurons may naturally be more likely to be remembered. Another possibility is that IT representations are explicitly shaped to prioritize memorability, reflecting evolutionary pressures to retain certain kinds of visual information. Our results cannot fully distinguish between these accounts, but they provide evidence for the latter by showing that memorability variance is complementary to recognition variance and can be learned as a separable optimization target. In support of this idea, Lin et al. ^41^ propose that images that are harder to reconstruct from high-level visual features leave stronger memory traces, suggesting that the interface between perception and memory is governed by the efficiency of perceptual encoding rather than memorability per se. Their sparse-coding model shows that reconstruction error—a signal derived from compressing and reconstructing deep network activations—predicts human memorability beyond standard vision-only measures. This framework provides a mechanistic account for how perceptual processing depth could modulate IT activity and, in turn, memory encoding, complementing our finding that memorability variance forms an explicit, learnable component of IT representations.

Functionally, memorability may serve as a prioritization signal for downstream memory systems, like the hippocampal formation^42^. By amplifying representations of images that are intrinsically memorable, IT may bias hippocampal encoding toward information that is more likely to be useful in future behavior. This perspective is consistent with theories of predictive^43^ and goal-driven^26^ coding, in which sensory representations are optimized not just for accurate perception but also for guiding learning and decision-making.

### Broader relevance across domains

Our results also connect to broader themes in systems neuroscience. In auditory^44^ and motor domains^45^, representational spaces have been shown to reflect task goals and behavioral demand. In language, neural responses are better captured by models trained on both semantic and syntactic objectives than by models trained on either alone^46^. The present study contributes to this growing body of evidence, supporting the view that cortical representations typically arise from the integration of multiple representational goals.

For artificial intelligence (AI), the implication is clear: building brain-like models will require training on objectives that extend beyond recognition^47^. From a computational standpoint, multi-goal optimization may rely on mechanisms analogous to mixed selectivity in biological circuits, which increase representational dimensionality and support flexible behavior across diverse tasks^48^. Incorporating analogous mechanisms into ANNs could similarly enhance their ability to generalize across objectives while preserving brain-like structure. Memorability is one such dimension, but others—such as emotional salience^49^, or language production^50^—may provide additional axes of variance. Incorporating these objectives may yield models that not only predict neural responses more accurately but also generalize more robustly across tasks, echoing the multi-goal optimization strategies that appear to guide biological systems.

### Reverse Predictivity improvements under joint optimization

In this study, we observed that joint optimization improved both forward and reverse predictivity (**Figure 5**). Reverse predictivity — the ability to reconstruct model feature space from IT neural responses — captures aspects of brain–model alignment that forward predictivity alone cannot. In previous work^35^, improvements in forward predictivity (by increasing effective dimensionality^51^, increasing the number of model parameters etc.) typically came at the expense of reverse predictivity. This suggested that common optimization strategies yielded representational spaces that were predictive of IT but not reciprocally embedded within IT responses. Our findings here show that this tradeoff is not inherent. Recognition+memorability models exhibited significant gains in both forward and reverse explained variance, pointing to a training strategy that pushes networks toward representational geometries more deeply constrained by IT-like structure. Why might joint objectives have this effect? One possibility is that recognition and memorability exert complementary pressures: recognition enforces categorical separability, while memorability emphasizes distinctiveness within and across categories. Optimizing for both simultaneously may regularize internal representations, discouraging features that are highly diagnostic for recognition but idiosyncratic to the model, and instead favoring feature dimensions that generalize across both behavioral goals. Such regularization likely reduces non–brain-like variance and increases the overlap between neural and model feature spaces, yielding improvements in both forward (model-to-brain) and reverse (brain-to-model) predictivity. Another interpretation is that memorability serves as a constraint on representational richness. By rewarding networks for capturing dimensions that make images stand out in memory, joint training may force models to retain variance that aligns with broader IT population responses, rather than collapsing onto narrowly optimized recognition axes. This broader representational overlap explains why IT activity can not only be predicted more accurately from models but also used more effectively to reconstruct the model’s own representational space. Together, these results highlight that the forward–reverse tradeoff reported previously is not inevitable. Instead, the choice of training objectives can determine whether models converge onto representational spaces that align with IT in both directions. Our findings suggest that incorporating multiple behavioral constraints — here, recognition and memorability — may be essential for developing models whose internal geometry is reciprocally embedded in neural activity.

### Limitations and future directions

Several limitations of this study point to directions for future research. First, while memorability accounted for unique variance in IT responses, the effect size was much smaller than recognition. This suggests that memorability is only one of several pressures shaping IT and that additional objectives may be needed to capture the full richness of IT representations^38^. Future work should systematically examine other candidate goals, such as image typicality, aesthetic value, or task relevance. Second, our analyses focused on static image responses. Memorability may also be influenced by temporal dynamics and recurrent processing^22^, which are not captured by the feedforward architectures studied here. Incorporating time-resolved neural data and recurrent models^52,53^ may reveal whether memorability is encoded dynamically or emerges from interactions between IT and downstream memory systems. Third, while we demonstrated improvements in ANN–IT alignment, the gains were modest relative to the variance explained by recognition. Future studies should systematically explore^54^ scaling effects, such as whether larger models or alternative architectures (e.g., transformers) exhibit greater benefits from multi-goal optimization. Finally, our behavioral tasks were limited to binary object recognition and single-shot memorability judgments. More naturalistic paradigms involving continuous memory demands, visual search, or long-term learning may better reveal how memorability interacts with recognition in real-world settings. A complementary approach is to investigate effects of perturbed or compromised IT function (e.g., via pharmacological means or lesions) on image recognition versus memorability. Such studies, along with examining extra-IT contributions in regions like the medial temporal lobe, would inform hierarchical models that integrate multiple brain regions in object recognition and memorability.

In sum, this study provides evidence that models of the ventral stream shaped by multiple representational goals develop more brain-like internal responses. Memorability constitutes a complementary organizing principle alongside recognition and incorporating memorability into ANN training yields models that better capture IT responses. These findings challenge the single-goal view of ventral stream function and argue for a shift toward multi-goal optimization in computational neuroscience. However, it remains possible that both object recognition and memorability draw upon a more fundamental representational process that determines the degree of IT engagement. By moving beyond recognition as the sole objective, we take a step toward building models that capture not only “what we see” but also “what we remember,” offering a richer account of the computations underlying visual cognition.

## Methods

### Human subjects

We collected large-scale psychophysical data from 104 subjects using Amazon Mechanical Turk (MTurk). Participants completed the tasks on MTurk, an online crowdsourcing platform, for a payment of $15 CAD/hour. This experimental protocol involving human participants was approved by and in concordance with the guidelines of the York University Human Participants Review Subcommittee. Humans did not receive any additional training on the task. For the object discrimination task, to ensure high-quality behavioral data, we rejected all participants who performed below an accuracy of 0.6 (chance-level = 0.5) on the first 30 trials during each session. Furthermore, we evaluated the reliability of the image-level behavioral metrics as a function of the number of trials and found that the reliability of the mean image-level accuracy pattern increased with more repetitions per image and reached high values (∼0.8), nearly asymptoting with around 60 trials. For the memorability task (see below), we had catch trials. Participants who failed to detect any catch repetitions were flagged as low performing and excluded from analysis.

### Non-human primates

We have three adult male rhesus macaques (Macaca mulatta) as research subjects in our experiments. All data were collected, and animal procedures were performed, in accordance with the NIH guidelines, the Massachusetts Institute of Technology Committee on Animal Care, and the guidelines of the Canadian Council on Animal Care on the use of laboratory animals, and were also approved by the York University Animal Care Committee. Some of the neural data used in this study have been utilized in previous publications^32,33^.

### Visual stimuli

We used a combination of grayscale naturalistic images and colored photographs from the MS COCO dataset, which was previously used in a study. The 640 synthetic grayscale naturalistic images were rendered from a 3D object model (originally purchased from TurboSquid) into a 2D projection while varying the object’s position (horizontal and vertical), rotation (in the horizontal, vertical, or depth plane), and size. The rendered object view was then added to a randomly chosen natural image background, consisting of both indoor and outdoor scenes obtained from Dosch Design (www.doschdesign.com). All resulting images were gray-scaled and had a resolution of 256×256 pixels. The images contained 8 objects (bear, elephant, face, apple, car, dog, chair, plane), and 80 images per object. The MS COCO images were colored and also contained the object types from the 8 categories mentioned above, as well as birds and zebras (making a total of 10 object types, with 20 images per object type).

### Behavioral Tasks

#### Object Discrimination Task

We measured human image object discriminability (88 subjects) using a binary object discrimination paradigm implemented on Amazon Mechanical Turk (MTurk) with *jsPsych* (version 6). Each trial started with a 100-ms presentation of the sample image (1 out of 1,320 images). This was followed by a blank gray screen for 100 ms followed by a choice screen with the target and distractor objects. The subjects indicated their choice by touching the screen or clicking the mouse over the target object. Each subject saw an image only once. We collected the data such that there were 80 unique subject responses per image with varied distractor objects.

#### Image Memorability Task

We measured human image memorability using a continuous recognition paradigm implemented on Amazon Mechanical Turk (MTurk) with *jsPsych* (version 6). Each Human Intelligence Task (HIT) began with an online consent form, followed by demographic questions (age, sex, education), task instructions, and full-screen enforcement to standardize stimulus presentation. On each trial, participants fixated on a central cross (500 ms), after which an image was shown for 100 ms and then replaced by a recognition prompt. Participants were instructed to indicate whether the image was “novel” (first presentation) or “repeated” (previously seen) using on-screen buttons, with a maximum response window of 2 seconds.

The experiment comprised five unique sessions, each containing 50 trials, and each session was repeated 22 times. Within each session, 10 trials were *target images* (50 unique targets across all sessions), which were presented twice: once as novel and once as repeated after a fixed lag of 10 intervening trials. The remaining 40 trials consisted of filler images, 4 of which served as “catch” trials that were repeated immediately (lag = 0) to monitor attention. Participants who failed to detect any catch repetitions were flagged as low performing and excluded from analysis.

Image size was scaled dynamically according to the participant’s screen resolution to subtend ∼8° of visual angle. A progress bar was displayed throughout the task to encourage compliance. All responses and reaction times were logged automatically by *jsPsych* and submitted to MTurk upon task completion.

#### Passive fixation

During the passive viewing task, monkeys fixated on a white circle (0.2°) for 300 ms to initiate a trial. We then presented a sequence of 5–10 images, each one for 100 ms, followed by a 100 ms gray (background) blank screen. This was followed by a fluid reward and an inter-trial interval of 500 ms, followed by the next sequence. Trials were aborted if the gaze was not held within ±2° of the central fixation circle at any point. The 640 naturalistic, grayscale images were used for the passive fixation task.

#### Eye Tracking

Eye movements were monitored using an EyeLink 1000 video eye-tracking system (SR Research). Through operant conditioning, subjects were trained to fixate on a central 0.2° white circle within a ±2° fixation window. Each behavioral trial commenced with an eye calibration procedure, where the monkeys made saccades to spatial targets and maintained fixation for 500ms. Calibration was repeated if any drift was observed.

#### Neural Recordings and Inclusion Criteria

We surgically implanted each monkey with a head post under aseptic conditions. We recorded neural activity using two or three micro-electrode arrays (Utah arrays; Blackrock Microsystems) implanted in the IT cortex. A total of 96 electrodes were connected per array (grid arrangement, 400 um spacing, 4mm × 4mm span of each array). Array placement was guided by the sulcus pattern, which was visible during the surgery. The electrodes were accessed through a percutaneous connector that allowed simultaneous recording from all 96 electrodes from each array. All data were collected, and animal procedures were performed, in accordance with the NIH guidelines, the Massachusetts Institute of Technology Committee on Animal Care, and the guidelines of the Canadian Council on Animal Care on the use of laboratory animals and were also approved by the York University Animal Care Committee.

During each daily recording session, band-pass filtered (0.1 Hz to 10 kHz) neural activity was recorded continuously at a sampling rate of 20 kHz using Intan Recording Controllers (Intan Technologies, LLC). The majority of the data presented here were based on multiunit activity. We detected the multiunit spikes after the raw voltage data were collected. A multiunit spike event was defined as the threshold crossing when voltage (falling edge) deviated by less than three times the standard deviation of the raw voltage values. Our array placements allowed us to sample neural sites from different parts of IT, along the posterior to anterior axis. However, for all the analyses, we did not consider the specific spatial location of the site and treated each site as a random sample from a pooled IT population.

#### Neural site inclusion criteria

For our analyses, we only included neural recording sites that exhibited an overall significant visual drive, an image rank order response reliability greater than 0.7Given that most of our neural metrics are corrected for the estimated noise at each neural site, the criterion for selecting neural sites is not critical, and it was primarily used to reduce computation time by eliminating noisy recordings.

To assess the reliability of individual neural sites, we computed a split-half internal consistency metric across stimulus repetitions. This measure quantifies how stable each site’s response pattern is across random subsets of trials, providing an inclusion criterion for downstream analyses (e.g., only sites exceeding a reliability threshold of 0.7 were used for model–neural comparisons).

For each neural site, we repeatedly divided the available trials into two random, non-overlapping halves. Within each split, we computed the mean response of the neuron across all repetitions for each image. The correlation between these two split means was taken as the raw split-half reliability. To obtain a reliability estimate corrected for finite sampling, we applied the Spearman–Brown correction, which compensates for the halving of the data. This procedure was repeated across 100 random half-splits, and the final reliability for each site was defined as the average of these corrected correlations.

### ANN model analyses

#### Overview

We asked whether image memorability contributes a signal complementary to object recognition, and whether combining both objectives improves model alignment with downstream targets. We trained a panel of 26 convolutional and transformer-based architectures under three regimes: (i) memorability-only (trained on LaMem dataset), (ii) distillation (Learning-without-Forgetting, LwF) to retain recognition while learning memorability, and (iii) random-label controls with matched sample counts that ablate the image–memorability signal. We instantiated 26 off-the-shelf image classification backbones spanning modern CNNs and Vision Transformers (e.g., VGG, ResNet, EfficientNet, ConvNeXt, ViT, Swin families; full list in Table 1). Each backbone fed two lightweight task heads: (1) a classification head (used only for distillation/recognition retention) and (2) a single-unit regression head for memorability.

#### Backbone initialization and trainability

The backbone is the feature extractor preceding the task-specific heads. In the memorability-only regime, models were initialized from random weights, and the backbone was fully trainable, optimized on LaMem using the regression head. In the distillation routine^34^, teacher and student backbones were initialized from ImageNet pretraining; the teacher was frozen to provide soft targets, while the student remained trainable and was updated under a combined objective comprising the memorability regression loss and the distillation loss.

### Datasets and preprocessing

#### Memorability (using the LaMem dataset^30^)

We trained on LaMem images with scalar memorability scores using standard preprocessing (resize to the model’s nominal input size, tensor conversion, and normalization with ImageNet statistics). Unless otherwise specified, no additional data augmentation was applied.

#### Object recognition (using the ImageNet dataset^14^)

For object recognition only models, we downloaded the pretrained models from PyTorch model zoo. For the joint optimized models, to monitor recognition retention in distillation runs, the student was evaluated on the ILSVRC-2012^68^ validation set using the conventional preprocessing pipeline; for efficiency, evaluation used a fixed subset of validation batches each epoch (subset size specified a priori).

### Model Training

#### Memorability-only (from scratch)

For each architecture, we trained from scratch on LaMem with an MSE loss on the 1-unit regression head. Optimization used Adam (learning rate 1 × 10 ^−3^, batch size 32, 30 epochs; betas *β*_1_ =0.9, *β*_2_ =0.999), with StepLR (step =10 epochs, *γ* = 0.1); weight decay was 0.0.

#### Distillation to retain recognition while learning memorability (LwF)

We employed a teacher–student protocol consistent with knowledge distillation and Learning-without-Forgetting^34^. For each LaMem minibatch, the student optimized a memorability regression loss together with a distillation loss on the classification head using temperature-softened teacher targets; the total objective was a convex combination with *T* = 2.0 and *α* = 0.4. Optimization used Adam (learning rate 1 × 10 ^−4^, batch size 32, 30 epochs; betas as above) with StepLR (*step* = 10, *γ* = 0.1) and *W* = 0.0. Recognition retention was monitored via Top-1/Top-5 accuracy of the student on a fixed subset of ImageNet validation batches each epoch; all parameter updates were driven exclusively by LaMem images (no ImageNet training).

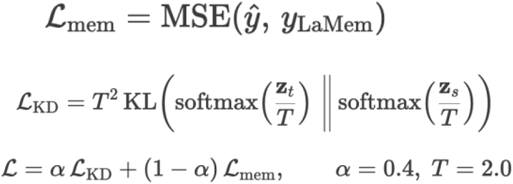

#### Random labels (negative-control training)

As a control analysis, we replaced memorability scores with random labels independent of both images and their corresponding true memorability. For each LaMem image, we sampled *u ∼* Uniform[0,1] with a fixed base seed and repeatedly re-sampled until the held-out Pearson correlation with true memorability was approximately zero (statistically indistinguishable from zero). This control was run under both regimes—memorability-only (from scratch) and distillation (LwF)—with all other data, hyperparameters, and schedules held constant. By design, the random-label condition exhibits near-zero correlation with true memorability, isolating effects attributable to the memorability signal from those due to additional optimization, regularization, or knowledge distillation itself.

#### Implementation details (reproducibility)

Experiments were implemented in PyTorch and executed on NVIDIA A100 GPUs. We used fixed random seeds for initialization and data shuffling; checkpoints and logs were saved each epoch. Unless otherwise stated, optimization settings and schedules followed those specified above.

### Prediction of neural activity from ANN features and vice versa

#### Forward Predictivity Analysis

To quantify forward predictivity, we evaluated how well each model’s features could linearly predict IT neural responses, following standard procedures. For each model and each recorded neuron *j*, we fit a linear regression (ridge regression) to predict the neuron’s responses across images from the model’s activations, with a cross-validation procedure (see Cross-Validation section). Given a response vector *r*_*j*_ ∈^1×*N*^ and a model activation matrix *A* ∈^*M*×*N*^, we estimated weights *w*_*j*_ ∈^*M*×1^ and bias *b* to minimize the squared error:

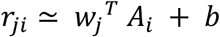

where *A*_*i*_ is the model feature vector for a given image *i*.

Model performance was assessed using explained variance (EV), defined as the squared Pearson correlation between predicted and actual firing rates, normalized by the geometric mean of their split-half reliabilities, and scaled to percentage. We computed %EV for each neuron, of each monkey and for each model. A model’s overall forward predictivity score was defined as the mean %EV across the full population of neurons.

### Reverse Predictivity Analysis

Reverse predictivity measures how well the recorded IT neuron population can predict the activations of individual units within each model’s IT-layer representation. For each model unit *k*, we treated its responses across images as the target variable and used the corresponding neural responses as predictors. Specifically, we fit a linear model of the form:

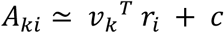

where *A*_*ki*_ is the activation of a model unit *k* for image *i, r*_*i*_ is the vector of IT neural responses for that image, *v*_*k*_ is a weight vector, and *c* is a bias term. We used linear regression (ridge regression) with cross-validation to fit the model for each unit. Predictive performance was quantified using explained variance (%EV), computed as the squared Pearson correlation between predicted and actual activations, normalized by the geometric mean of the split-half reliabilities of the prediction and the target, and expressed as a percentage.

We computed EV for each model unit predicted from each monkey IT cortex. A model’s overall reverse predictivity score was defined as the mean %EV across all units.

### Representational similarity analysis (CKA)

In centered kernel alignment (CKA^31^), the alignment between the activation features of different layers is measured by comparing the inner dot products. This technique utilizes the Hilbert-Schmidt Independence Criterion to quantify the similarity between the activation features of layers, considering both their linear and non-linear relationships.

### Statistical Analyses

The Shapiro-Wilk Test was employed to determine the normality of the data distribution. Based on the results of these tests, the appropriate use of parametric and non-parametric tests was determined. For normally distributed data with equal variances, parametric tests such as the one-sample T-test and the two-sample T-test were used. The non-parametric Mann-Whitney U test was used as an alternative when the assumptions of normality or homoscedasticity were not met.

#### Correlational analyses

Partial correlation analysis was conducted to determine if memorability and recognition models explain unique or overlapping variance in the neural predictions. In partial correlation, the relationship between two variables is examined using Spearman’s correlation while controlling for the influence of a third variable. In all other instances of correlational analyses, the type of correlation used, the correlation values, and the p-values have been reported.

## Acknowledgments

KK has been supported by funds from the Canada Research Chair Program (CRC-2021-00326), the Canada First Research Excellence Funds (VISTA Program), and the National Sciences and Engineering Research Council of Canada (NSERC, RGPIN-2024-06223). SM is funded by the Connected Minds Postdoctoral Fellowship (supported by CFREF). EF is supported by the Translating Brain Signal Grant (York University) and the VISTA Program (supported by CFREF). RSR acknowledges support from a York Research Chair, NSERC, and the CFREF VISTA and Connected Minds programs.

## Supplementary Material

**Figure S1.**
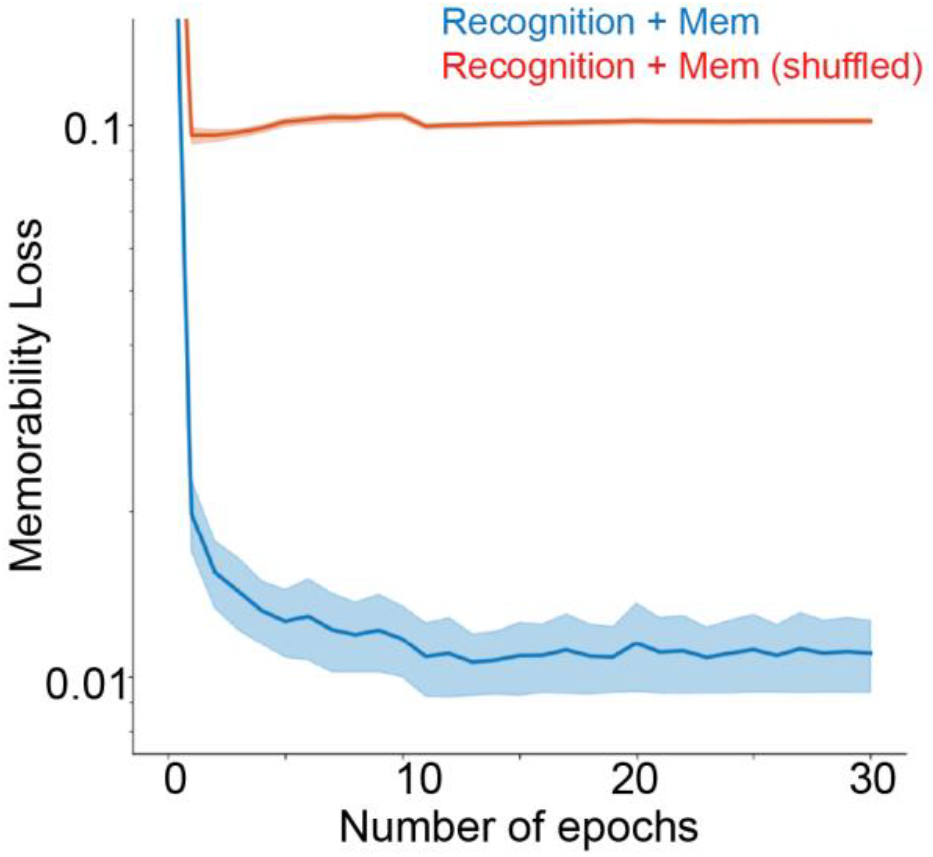
Memorability loss decreases only when trained with true memorability labels. Training curves for models jointly optimized for recognition and memorability (blue) versus recognition with shuffled memorability labels (red). Memorability loss rapidly decreased and stabilized over training epochs only when models were trained with true memorability scores, whereas loss remained high and unchanged when memorability labels were randomized. Shaded area represents ±1 SEM across architectures.

**Figure S2.**
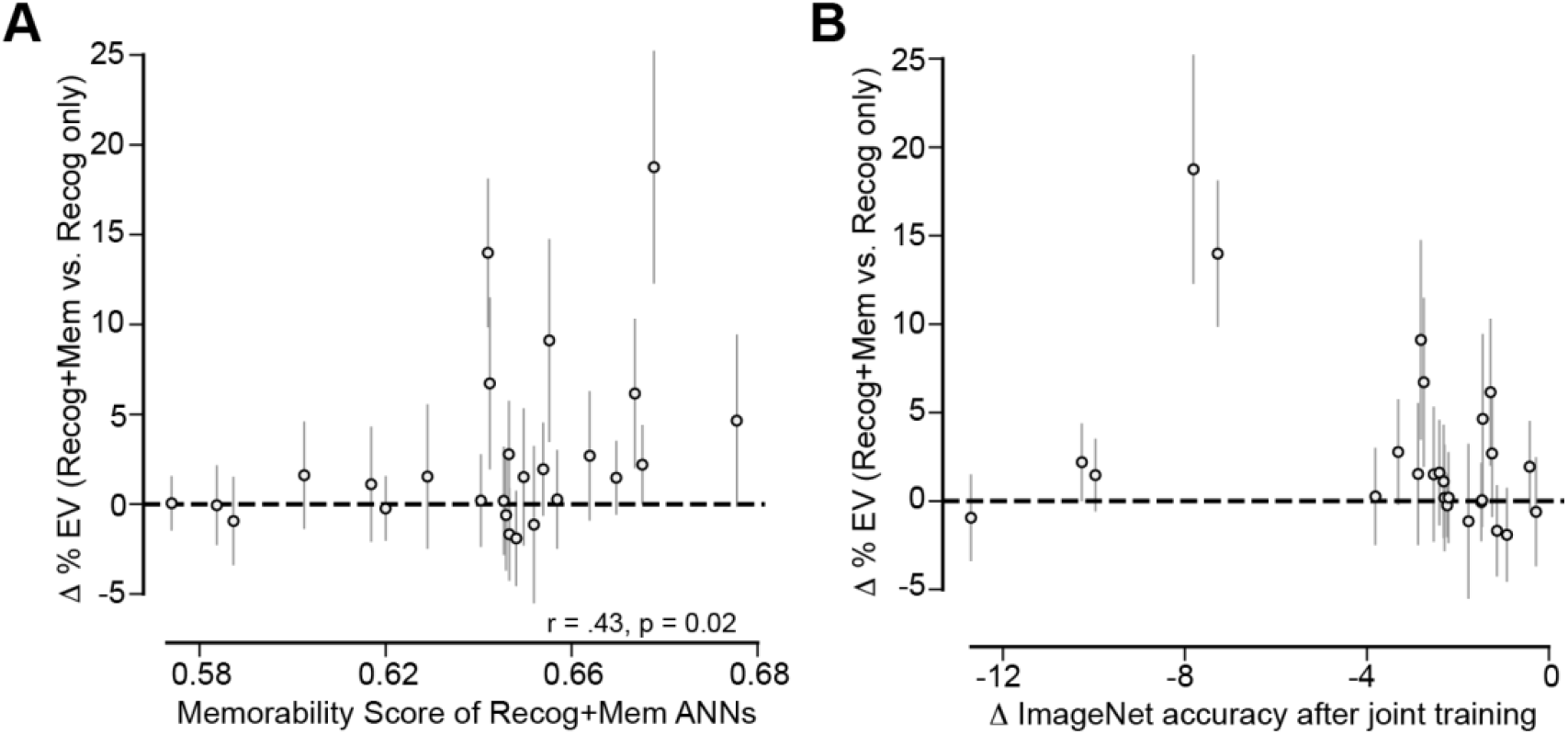
Gains in IT predictivity from joint optimization scale with memorability performance and depend on moderate changes in ImageNet accuracy. A. Improvement in IT predictivity (Δ%EV) between jointly optimized (Recognition + Memorability) and recognition-only models plotted against the memorability scores of the joint models. Models that achieved higher memorability performance showed greater increases in IT explained variance (r = 0.43, *p* = 0.02). B. The same measure plotted against the change in ImageNet classification accuracy (Δ accuracy) after joint training. Gains in IT alignment were largest for models showing moderate reductions in ImageNet accuracy, whereas models with minimal or large decreases in accuracy exhibited little or no change in %EV. This pattern indicates that modest adjustments in recognition performance are most conducive to capturing memorability- related variance in IT. Error bars denote ±1 s.e.m. across neurons.

**Figure S3.**
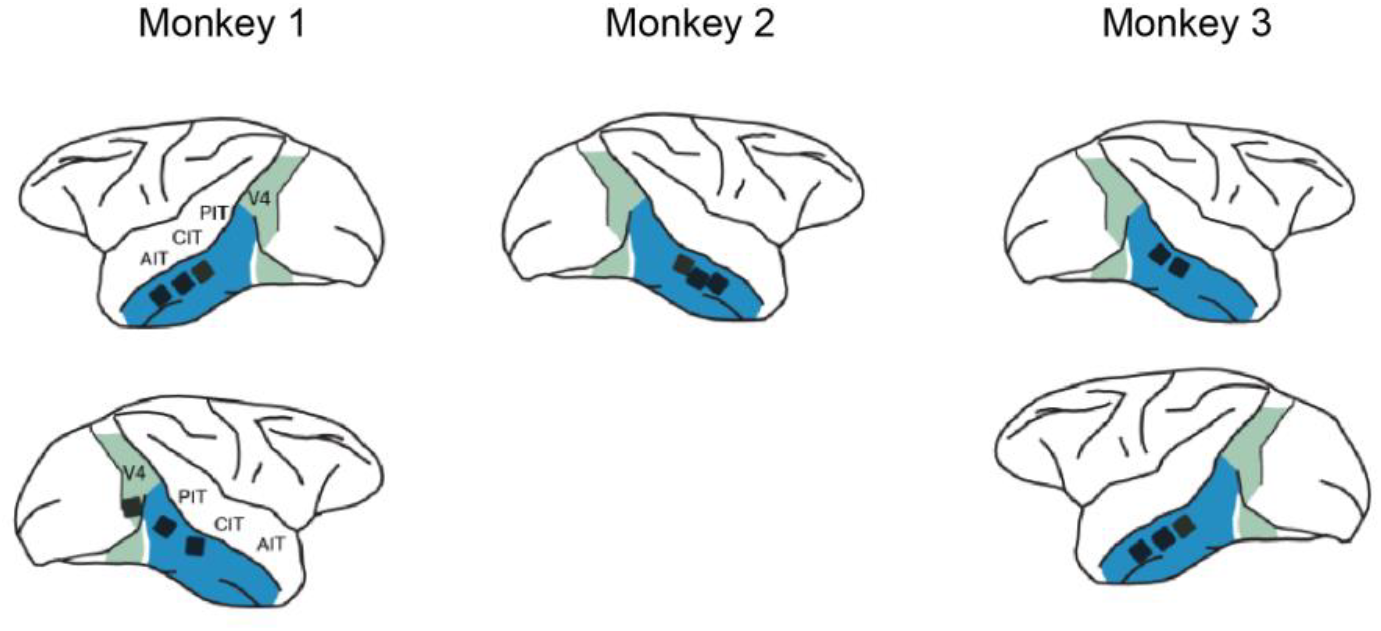
Schematic of Utah array placement in the three monkeys used in the study.

**Figure S4.**
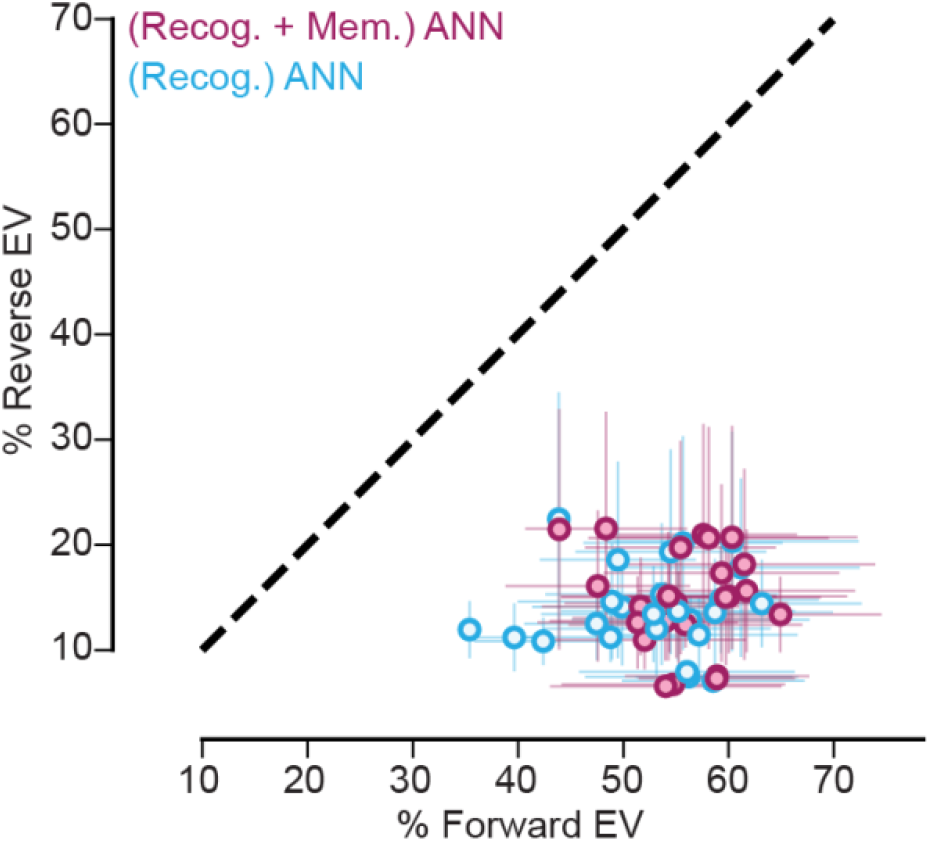
Forward vs. Reverse Predictivity across all 26 model architectures (trained on recognition only – light blue – and jointly optimized with memorability – purple). In both cases, forward predictivity %EV (models predict IT responses) is higher than reverse predictivity %EV (IT responses predict model units). However, the joint optimization pushes the dots along both axes, increasing scores in both forward and reverse predictivities. Each dot represents the mean %EV with error bars indicating median absolute deviation (m.a.d.) across neurons/units.

## Notes

**Conflict of interest** The author declares no competing financial interests.

https://github.com/vital-kolab/ann-augment-memorability

### Competing Interest Statement

The authors have declared no competing interest.

## References

1. DiCarlo, J. J., Zoccolan, D. & Rust, N. C. How Does the Brain Solve Visual Object Recognition? Neuron 73, 415–434 (2012).

2. Kar, K. & DiCarlo, J. J. The Quest for an Integrated Set of Neural Mechanisms Underlying Object Recognition in Primates. Annu. Rev. Vis. Sci. 10, 91–121 (2024).

3. Hung, C. P., Kreiman, G., Poggio, T. & DiCarlo, J. J. Fast Readout of Object Identity from Macaque Inferior Temporal Cortex. Science 310, 863–866 (2005).

4. Logothetis, N. K. & Sheinberg, D. L. Visual Object Recognition. Annu. Rev. Neurosci. 19, 577–621 (1996).

5. Tsao, D. Y., Schweers, N., Moeller, S. & Freiwald, W. A. Patches of face-selective cortex in the macaque frontal lobe. Nat. Neurosci. 11, 877–879 (2008).

6. Afraz, A., Boyden, E. S. & DiCarlo, J. J. Optogenetic and pharmacological suppression of spatial clusters of face neurons reveal their causal role in face gender discrimination. Proc. Natl. Acad. Sci. 112, 6730–6735 (2015).

7. Afraz, S.-R., Kiani, R. & Esteky, H. Microstimulation of inferotemporal cortex influences face categorization. Nature 442, 692–695 (2006).

8. Rajalingham, R. & DiCarlo, J. J. Reversible Inactivation of Different Millimeter-Scale Regions of Primate IT Results in Different Patterns of Core Object Recognition Deficits. Neuron 102, 493-505.e5 (2019).

9. James, T. W., Culham, J., Humphrey, G. K., Milner, A. D. & Goodale, M. A. Ventral occipital lesions impair object recognition but not object-directed grasping: an fMRI study. Brain 126, 2463–2475 (2003).

10. Liu, T. T. & Behrmann, M. Functional outcomes following lesions in visual cortex: Implications for plasticity of high-level vision. Neuropsychologia 105, 197–214 (2017).

11. DiCarlo, J. J. & Cox, D. D. Untangling invariant object recognition. Trends Cogn. Sci. 11, 333–341 (2007).

12. Yamins, D. L. K. et al. Performance-optimized hierarchical models predict neural responses in higher visual cortex. Proc. Natl. Acad. Sci. 111, 8619–8624 (2014).

13. Schrimpf, M. et al. Brain-Score: Which Artificial Neural Network for Object Recognition Is Most Brain-Like? http://biorxiv.org/lookup/doi/10.1101/407007 (2018) xdoi:10.1101/407007.

14. Deng, J. et al. ImageNet: A large-scale hierarchical image database. in 2009 IEEE Conference on Computer Vision and Pattern Recognition 248–255 (2009). doi:10.1109/CVPR.2009.5206848.

15. Cadieu, C. F. et al. Deep Neural Networks Rival the Representation of Primate IT Cortex for Core Visual Object Recognition. PLoS Comput. Biol. 10, e1003963 (2014).

16. Rajalingham, R. et al. Large-Scale, High-Resolution Comparison of the Core Visual Object Recognition Behavior of Humans, Monkeys, and State-of-the-Art Deep Artificial Neural Networks. J. Neurosci. 38, 7255–7269 (2018).

17. Khaligh-Razavi, S.-M. & Kriegeskorte, N. Deep Supervised, but Not Unsupervised, Models May Explain IT Cortical Representation. PLoS Comput. Biol. 10, e1003915 (2014).

18. Hong, H., Yamins, D. L. K., Majaj, N. J. & DiCarlo, J. J. Explicit information for category-orthogonal object properties increases along the ventral stream. Nat. Neurosci. 19, 613–622 (2016).

19. Jaegle, A. et al. Population response magnitude variation in inferotemporal cortex predicts image memorability. eLife 8, e47596 (2019).

20. Ramezanpour, H., Ilic, F., Wildes, R. P. & Kar, K. Object motion representation in the macaque ventral stream – a gateway to understanding the brain’s intuitive physics engine. Preprint at 10.1101/2024.02.23.581841 (2024).

21. Rust, N. C. & Mehrpour, V. Understanding Image Memorability. Trends Cogn. Sci. 24, 557–568 (2020).

22. Lahner, B., Mohsenzadeh, Y., Mullin, C. & Oliva, A. Visual perception of highlymemorable images is mediated by a distributed network of ventral visual regions that enable a late memorability response. PLOS Biol. 22, e3002564 (2024).

23. Kramer, M. A., Hebart, M. N., Baker, C. I. & Bainbridge, W. A. The features underlying the memorability of objects. Sci. Adv. 9, eadd2981 (2023).

24. Bainbridge, W. A., Dilks, D. D. & Oliva, A. Memorability: A stimulus-driven perceptual neural signature distinctive from memory. NeuroImage 149, 141–152 (2017).

25. Richards, B. A. et al. A deep learning framework for neuroscience. Nat. Neurosci. 22, 1761–1770 (2019).

26. Yamins, D. L. K. & DiCarlo, J. J. Using goal-driven deep learning models to understand sensory cortex. Nat. Neurosci. 19, 356–365 (2016).

27. Kar, K., Kubilius, J., Schmidt, K., Issa, E. B. & DiCarlo, J. J. Evidence that recurrent circuits are critical to the ventral stream’s execution of core object recognition behavior. Nat. Neurosci. 22, 974–983 (2019).

28. Majaj, N. J., Hong, H., Solomon, E. A. & DiCarlo, J. J. Simple Learned Weighted Sums of Inferior Temporal Neuronal Firing Rates Accurately Predict Human Core Object Recognition Performance. J. Neurosci. 35, 13402–13418 (2015).

29. Djambazovska, S., Zafer, A., Ramezanpour, H., Kreiman, G. & Kar, K. The Impact of Scene Context on Visual Object Recognition: Comparing Humans, Monkeys, and Computational Models. BioRxiv Prepr. Serv. Biol. 2024.05.27.596127 (2024) doi:10.1101/2024.05.27.596127.

30. Khosla, A., Raju, A. S., Torralba, A. & Oliva, A. Understanding and Predicting Image Memorability at a Large Scale. in 2015 IEEE International Conference on Computer Vision (ICCV) 2390–2398 (IEEE, Santiago, Chile, 2015). doi:10.1109/ICCV.2015.275.

31. Kornblith, S., Norouzi, M., Lee, H. & Hinton, G. Similarity of Neural Network Representations Revisited. Preprint at 10.48550/arXiv.1905.00414 (2019).

32. Sörensen, L. K. A., DiCarlo, J. J. & Kar, K. The effects of object category training on the responses of macaque inferior temporal cortex are consistent with performance-optimizing updates within a visual hierarchy. Preprint at 10.1101/2024.12.27.630539 (2024).

33. Dapello, J. et al. Aligning Model and Macaque Inferior Temporal Cortex Representations Improves Model-to-Human Behavioral Alignment and Adversarial Robustness. in The Eleventh International Conference on Learning Representations (2022).

34. Li, Z. & Hoiem, D. Learning without Forgetting. Preprint at 10.48550/arXiv.1606.09282 (2017).

35. Muzellec, S. & Kar, K. Reverse Predictivity: Going Beyond One-Way Mapping to Compare Artificial Neural Network Models and Brains. Preprint at 10.1101/2025.08.08.669382 (2025).

36. Geirhos, R. et al. ImageNet-trained CNNs are biased towards texture; increasing shape bias improves accuracy and robustness. ArXiv181112231 Cs Q-Bio Stat http://arxiv.org/abs/1811.12231 (2019).

37. Geirhos, R. et al. Generalisation in humans and deep neural networks. ArXiv180808750 Cs Q-Bio Stat http://arxiv.org/abs/1808.08750 (2020).

38. Xie, Y. et al. Vision CNNs trained to estimate spatial latents learned similar ventral-stream-aligned representations. Preprint at 10.48550/arXiv.2412.09115 (2025).

39. Kanwisher, N., Khosla, M. & Dobs, K. Using artificial neural networks to ask ‘why’ questions of minds and brains. Trends Neurosci. 46, 240–254 (2023).

40. Tuckute, G. et al. How to optimize neuroscience data utilization and experiment design for advancing brain models of visual and linguistic cognition? Preprint at 10.48550/ARXIV.2401.03376 (2024).

41. Lin, Q., Li, Z., Lafferty, J. & Yildirim, I. Images with harder-to-reconstruct visual representations leave stronger memory traces. Nat. Hum. Behav. 8, 1309–1320 (2024).

42. Voss, J. L., Bridge, D. J., Cohen, N. J. & Walker, J. A. A Closer Look at the Hippocampus and Memory. Trends Cogn. Sci. 21, 577–588 (2017).

43. Rao, R. P. & Ballard, D. H. Predictive coding in the visual cortex: a functional interpretation of some extra-classical receptive-field effects. Nat. Neurosci. 2, 79–87 (1999).

44. Kell, A. J. E., Yamins, D. L. K., Shook, E. N., Norman-Haignere, S. V. & McDermott, J. H. A Task-Optimized Neural Network Replicates Human Auditory Behavior, Predicts Brain Responses, and Reveals a Cortical Processing Hierarchy. Neuron 98, 630-644.e16 (2018).

45. Michaels, J. A., Schaffelhofer, S., Agudelo-Toro, A. & Scherberger, H. A goal-driven modular neural network predicts parietofrontal neural dynamics during grasping. Proc. Natl. Acad. Sci. 117, 32124–32135 (2020).

46. Schrimpf, M. et al. The neural architecture of language: Integrative modeling converges on predictive processing. Proc. Natl. Acad. Sci. 118, e2105646118 (2021).

47. Yang, G. R., Joglekar, M. R., Song, H. F., Newsome, W. T. & Wang, X.-J. Task representations in neural networks trained to perform many cognitive tasks. Nat. Neurosci. 22, 297–306 (2019).

48. Fusi, S., Miller, E. K. & Rigotti, M. Why neurons mix: high dimensionality for higher cognition. Curr. Opin. Neurobiol. 37, 66–74 (2016).

49. Kragel, P. A., Reddan, M. C., LaBar, K. S. & Wager, T. D. Emotion schemas are embedded in the human visual system. Sci. Adv. 5, eaaw4358 (2019).

50. Doerig, A. et al. High-level visual representations in the human brain are aligned with large language models. Nat. Mach. Intell. 7, 1220–1234 (2025).

51. Elmoznino, E. & Bonner, M. F. High-performing neural network models of visual cortex benefit from high latent dimensionality. PLOS Comput. Biol. 20, e1011792 (2024).

52. Nayebi, A. et al. Recurrent Connections in the Primate Ventral Visual Stream Mediate a Trade-Off Between Task Performance and Network Size During Core Object Recognition. Neural Comput. 34, 1652–1675 (2022).

53. Kubilius, J. et al. Brain-Like Object Recognition with High-Performing Shallow Recurrent ANNs. in Advances in Neural Information Processing Systems (eds Wallach, H.et al.) vol. 32 (Curran Associates, Inc., 2019).

54. Conwell, C., Prince, J. S., Kay, K. N., Alvarez, G. A. & Konkle, T. A large-scale examination of inductive biases shaping high-level visual representation in brains and machines. Nat. Commun. 15, 9383 (2024).

55. Krizhevsky, A., Sutskever, I. & Hinton, G. E. ImageNet Classification with Deep Convolutional Neural Networks. in Advances in Neural Information Processing Systems (eds Pereira, F., Burges, C. J. C., Bottou, L. & Weinberger, K.Q.) vol. 25 (Curran Associates, Inc., 2012).

56. He, K., Zhang, X., Ren, S. & Sun, J. Deep Residual Learning for Image Recognition. Preprint at http://arxiv.org/abs/1512.03385 (2015).

57. Simonyan, K. & Zisserman, A. Very Deep Convolutional Networks for Large-Scale Image Recognition. ArXiv14091556 Cs http://arxiv.org/abs/1409.1556 (2015).

58. Szegedy, C., Vanhoucke, V., Ioffe, S., Shlens, J. & Wojna, Z. Rethinking the Inception Architecture for Computer Vision. Preprint at 10.48550/arXiv.1512.00567 (2015).

59. Dosovitskiy, A. et al. An Image is Worth 16×16 Words: Transformers for Image Recognition at Scale. Preprint at 10.48550/arXiv.2010.11929 (2021).

60. Liu, Z. et al. Swin Transformer: Hierarchical Vision Transformer using Shifted Windows. Preprint at 10.48550/arXiv.2103.14030 (2021).

61. Iandola, F. N. et al. SqueezeNet: AlexNet-level accuracy with 50x fewer parameters and <0.5MB model size. Preprint at 10.48550/arXiv.1602.07360 (2016).

62. Ma, N., Zhang, X.Zheng, H.-T. & Sun, J. ShuffleNet V2: Practical Guidelines for Efficient CNN Architecture Design. Preprint at 10.48550/arXiv.1807.11164 (2018).

63. Sandler, M., Howard, A., Zhu, M., Zhmoginov, A. & Chen, L.-C. MobileNetV2: Inverted Residuals and Linear Bottlenecks. Preprint at 10.48550/arXiv.1801.04381 (2019).

64. Tan, M. & Le, Q. V. EfficientNetV2: Smaller Models and Faster Training. Preprint at 10.48550/arXiv.2104.00298 (2021).

65. Tan, M. & Le, Q. V. EfficientNet: Rethinking Model Scaling for Convolutional Neural Networks. Preprint at http://arxiv.org/abs/1905.11946 (2020).

66. Huang, G., Liu, Z., van der Maaten, L. & Weinberger, K. Q. Densely Connected Convolutional Networks. Preprint at 10.48550/arXiv.1608.06993 (2018).

67. Liu, Z. et al. A ConvNet for the 2020s. Preprint at 10.48550/arXiv.2201.03545 (2022).

68. Russakovsky, O. et al. ImageNet Large Scale Visual Recognition Challenge. Int. J. Comput. Vis. 115, 211–252 (2015).

